# From life history to proteome via mutagenesis: an ecological footprint in genome evolution

**DOI:** 10.64898/2026.08.14.744874

**Authors:** Alexandr Voronka, Anastasia Koshel, Gleb Osadchiy, Bogdan Efimenko, Konstantin Gunbin, Konstantin Popadin

**Author notes:** equal contribution (Alexandr Voronka, Anastasia Koshel, Gleb Osadchiy). equal contribution.

## Abstract

Species differ in longevity, physiology, and social organization, and these properties expose them to distinct endogenous and environmental mutagens. Mutational spectra generated by these processes can shape downstream molecular evolution, influencing synonymous nucleotide composition, codon usage, and even amino acid composition. Tracing this signal from life history to proteome through mutagenesis could reveal how mutational pressure interacts with the fitness landscape, including the direction of molecular change and the extent to which proteins remain functional while following mutational biases. Here, building on the recently identified age-associated mitochondrial A>G mutational signature in mammals, we test the universality of this signature and its downstream effects on genome and proteome evolution by comparing long-lived termites with short-lived non-termite cockroaches. We find that termite mtDNA exhibits a stronger A>G mutational signature than that of non-termite cockroaches, accompanied by coordinated shifts in synonymous nucleotide composition, codon usage, and amino acid composition. Our results show that ecological and life-history-associated mutational pressures can be transmitted through a hierarchy from mutational spectra to nucleotide composition and ultimately to proteome evolution. Mitochondrial genomes may therefore function not only as records of ancestry but also as molecular archives of the biological conditions under which species evolve.

## INTRODUCTION

Mitochondrial DNA (mtDNA) is vulnerable to asymmetric mutagenesis because one of its chains remains single-stranded for extended periods during strand-asynchronous replication. This prolonged single-stranded state exposes mtDNA to damage, including cytosine and adenine deamination, leading to C>T and A>G mutations [1]. Of these two common transitions in mtDNA, the A>G signature has also been associated with age-related damage in mammals [2], metabolism-associated damage in vertebrates [3], and temperature-associated damage in actinopterygians [4]. Together, these findings support the hypothesis that A>G substitutions on single-stranded mtDNA can serve as an informative marker of species-specific life-history traits, physiological properties, and ecological niches. Moreover, species-specific mutational spectra can drive downstream changes in synonymous nucleotide composition [2] and influence amino acid composition, as demonstrated both across diverse protein-coding sequences [5,6] and, more recently, through quantitative analyses of RNA-virus evolution [7].

In this study, we test the generality of the ecologically modulated mutagenesis hypothesis and its two downstream consequences by comparing closely related insects with markedly different longevities: termites and cockroaches. Although termites are phylogenetically nested within the cockroach lineage of Blattodea, they are characterized by exceptionally long-lived reproductive females, whereas most non-termite cockroaches are comparatively short-lived [8]. Hereafter, we use “cockroaches” to refer to non-termite cockroach lineages. These groups therefore provide an informative comparative system for testing how life history influences mtDNA evolution and whether long lifespan is associated with a stronger A>G-rich mitochondrial mutational signature, skewed synonymous nucleotide composition, and altered amino acid composition. We find a strong signature of age-associated mutagenesis in termite mtDNA, characterized by an elevated A>G rate on the strand that remains single-stranded, and show that this difference leaves a detectable imprint on both synonymous and amino acid-level mitochondrial evolution.

## RESULTS

### 1. Long-lived termites show a more A>G-rich mitochondrial mutational spectrum than short-lived cockroaches

We recently identified an age-related mutational signature in mammalian mtDNA, characterized by elevated A>G transitions on single-stranded DNA [2] (Fig. 1A). To test its universality across animal mtDNA, here we compared two insect groups with strikingly different lifespans: termites—whose extraordinarily long-lived queens far outlive other castes [8] —and shorter-lived cockroaches. Due to mtDNA’s exclusively maternal inheritance—which excludes contributions from other castes but queens —the extraordinarily long lifespan of termite queens is expected to directly shape its species-specific mtDNA mutagenesis and evolution. Using the NeMu tool [9] and MIDORI2 dataset [10] (see Methods), we derived COX-1-based mtDNA mutational spectra separately for each group (Fig. 1B). Among the 12 substitution types, A>G (hereafter we use the minority strand notation which correspond to heavy strand of vertebrate mtDNA) showed the strongest difference with termites exhibiting substantially higher frequencies (median 0.157, n=20) than cockroaches (median 0.074, n=20; Mann–Whitney p=0.00046). These results suggest the universality of the aging signature in mtDNA across distant taxa, implying that the underlying mutagens— single-stranded damage of adenine —can be similarly conserved.

**Figure 1.**
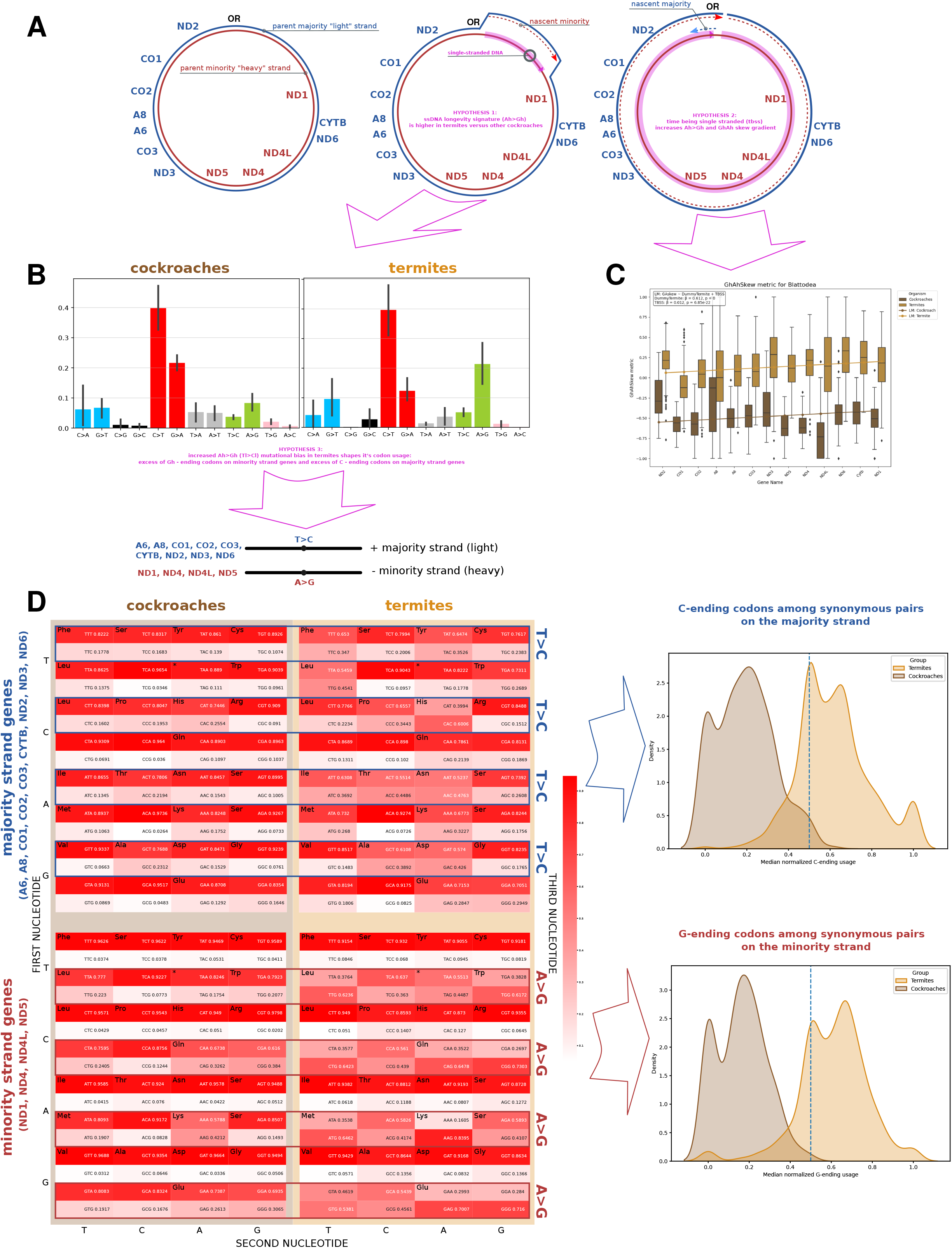
A termite-associated mitochondrial mutational bias propagates into nucleotide composition and codon usage. (high-quality image on **GitHub**) **A**, Model of strand-asynchronous mtDNA replication. The minority strand remains single-stranded for a prolonged period, providing a mechanistic basis for the A>G mutational bias and its predicted gradient along the mitochondrial genome. **B**, B, Twelve-component mitochondrial mutational spectra reconstructed from COX1 sequences of termite (N=20) and non-termite cockroach species (N=20). Substitutions are reported using minority-strand notation, reflecting the strand’s prolonged single-stranded exposure and predicted susceptibility to A>G damage. Termites show an elevated A>G frequency relative to non-termite cockroaches. **C**, GhAh skew, calculated as (G−A)/(G+A) across mitochondrial genes. GhAh skew is higher in termites than in cockroaches and increases in both groups with the time each gene spends in the single-stranded state during replication. **D**, Strand- and taxon-specific synonymous codon usage (331 termite species and 66 cockroach species). The termite-associated A>G bias is reflected in increased use of G-ending codons on minority-strand genes and C-ending codons on majority-strand genes.

### 2. Synonymous nucleotide content and codon usage reflect the stronger A>G mutational bias in termites

If synonymous sites evolve largely under mutational pressure, we expect the nucleotide content to mirror the mutational spectra (Fig. 1B), resulting in a stronger excess of G over A—captured by more positive values of the G-A skew metric (*G*−*A*)/(*G*+*A*)—in termites than in cockroaches. In line with our hypothesis we observed increased G-A skew in termites (Fig. 1C), interestingly, this skew is also correlated with gene position along the mtDNA (gene location on the X-axis), reflecting the expected time the minority strand spends in the single-stranded state during asymmetric replication. Such a positional trend is expected under asymmetric mtDNA replication, because genes replicated later remain single-stranded for longer. Although mtDNA replication modes in insects remain debated, the close resemblance of our results to those in mammals [2] supports asymmetric replication of mtDNA as a common mechanism in insects. As a consequence of this strong synonymous nucleotide skew, we observe a corresponding codon usage bias (A>G in the minority-strand genes and T>C in the majority-strand genes), which is stronger in termites than in cockroaches (Fig. 1D).

### 3. Amino-acid divergence follows the mutational spectrum

A>G mutations on the minority strand deplete adenine (Amin) and enrich guanine (Gmin), with corresponding thymine depletion (Tmaj) and cytosine enrichment (Cmaj) on the majority strand. If mutagenesis outweighs selection, this bias can drive amino acid shift [7]. To test it, we first focused on robust qualitative trends from first and second codon nucleotides: minority-strand genes are expected to become enriched in glycine and depleted in lysine and asparagine, while majority-strand genes are expected to become enriched in proline and depleted in phenylalanine and leucine (Fig. 2A). These effects are predicted to be stronger in termites than in cockroaches due to increased A>G rate (Fig 1). Comparing amino acid composition between termite and cockroach mtDNA-encoded genes, we observed the predicted pattern: termite minority-strand genes indeed were enriched in glycine and depleted in lysine and asparagine, whereas majority-strand genes were enriched in proline and depleted in phenylalanine and leucine (Fig. 2B). Accordingly, the predicted-gainers-to-losers ratios were consistently higher in termites across all individual genes (Fig. 2C), supporting mutational spectrum-driven amino acid shifts. Following these qualitative comparisons (Fig. 2A–C), we next quantitatively compared all observed and expected amino acid divergence between taxa (see methods and [7] for model details). Observed and expected differences were significantly positively correlated for genes encoded on both the minority and majority strands (Fig. 2D).

**Figure 2.**
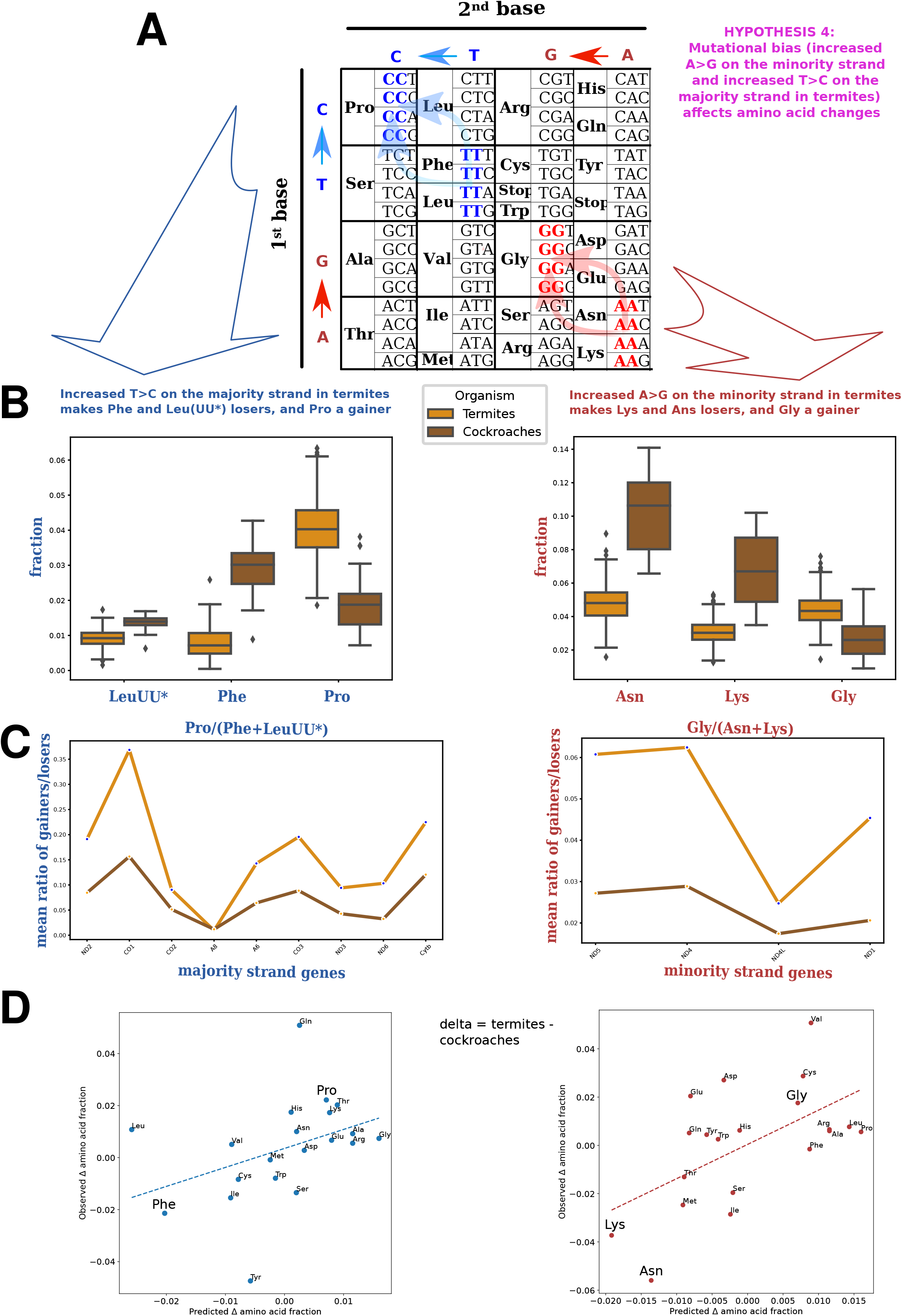
Mutational bias predicts amino acid composition in mitochondrial proteins (high-quality image on GitHub) **A**, Predicted amino acid consequences of the termite-associated mutational spectrum. Increased A>G mutagenesis in minority-strand genes is predicted to enrich glycine and deplete lysine and asparagine, whereas the corresponding T>C bias in majority-strand genes is predicted to enrich proline and deplete phenylalanine and leucine (TTX). **B**, Observed amino acid fractions in termite and cockroach mitochondrial genes, averaged separately across genes encoded on the minority and majority strands. Relative to cockroaches, termites show an excess of the predicted gainers (proline in majority-strand genes and glycine in minority-strand genes) and a deficit of the predicted losers (LeuTT and phenylalanine in majority-strand genes; asparagine and lysine in minority-strand genes). **C**, Ratio of predicted gainers to predicted losers for each mitochondrial gene. This ratio is consistently higher in termites than in cockroaches across both strands. **D**, Observed versus neutral-mutational-model-predicted amino acid differences between termites and cockroaches for majority- and minority-strand genes. The positive correlations support a substantial contribution of mutational bias to taxon-specific amino acid divergence.

## DISCUSSION

In this study, we find that termite mtDNA exhibits a stronger A>G mutational signature than that of non-termite cockroaches, and that coordinated shifts in synonymous nucleotide composition, codon usage, and amino acid composition accompany this difference. These results support a hierarchical model in which ecological, physiological, and life-history conditions influence mitochondrial mutagenesis, which in turn shapes nucleotide and protein evolution. Although positive selection can also be shaped by shifts in the mutational spectrum [11] [12] [13], the highly conserved mitochondrial genes examined here are unlikely to experience pervasive adaptive selection. The concordant changes in mutational spectra, fourfold-degenerate sites, codon usage, and amino acid composition are therefore most parsimoniously explained by a predominantly effectively neutral process, consistent with recent evidence from RNA viruses [7]. To estimate the contribution of mutational bias to amino acid evolution, we compared observed divergence across all mtDNA-encoded proteins of termites and cockroaches with divergence predicted from their mutational spectra (see Methods). Mutational bias was consistent with 78% of amino acid divergence in majority-strand genes and 92% in minority-strand genes, indicating a substantial contribution of mutagenesis to taxon-specific protein evolution. Although exploratory, these estimates demonstrate the strong predictive potential of mutational spectra.

This makes the result particularly notable: a mutation bias associated with organismal biology can propagate through the genome without requiring adaptive selection at every affected site. In this view, nucleotide and amino acid composition are not merely static properties of genomes, but molecular records of the mutational environments in which those genomes evolved. The direction of this relationship may even be reversed. If the relevant genomic regions and mutational signatures can be identified, sequence composition could provide a route for tracing ecological, physiological, and life-history characteristics from genomes—and potentially from proteins—back to the conditions that shaped their evolution.

The observation that this signal persists in the highly conserved mitochondrial protein-coding genome further suggests that even constrained proteins occupy sufficiently navigable fitness landscapes for mutational bias to influence their long-term evolutionary trajectories. Thus, the fitness landscape need not be traversed exclusively through positive selection: mutation pressure, drift, and weak selection may collectively redirect molecular evolution while preserving protein function. Our results therefore suggest that ecological information can be transmitted through a chain of predominantly natural events extending from organismal biology to mutational spectra, from mutational spectra to nucleotide composition, and from nucleotide composition to proteome evolution.

## MATERIALS AND METHODS

### 1. Data Collection

This study focuses on termites and cockroaches as closely related lineages within Blattodea that differ markedly in life-history traits and social organization. Species were selected to represent a broad taxonomic and ecological diversity within each group while ensuring sufficient availability and quality of mitochondrial sequence data. Termites included representatives of both lower and higher termites, whereas cockroaches encompassed primarily solitary species, with the inclusion of taxa exhibiting subsocial behavior. This sampling strategy allowed for comparative analyses across a gradient of social complexity within a shared evolutionary framework.

Mitochondrial protein-coding gene sequences were obtained from MIDORI database, which provides curated mitochondrial gene alignments linked to taxonomically validated species records. Only species for which complete or near-complete mitochondrial protein-coding gene coverage was available were considered. To reduce potential biases associated with sequencing errors or poor taxonomic resolution, sequences with ambiguous annotations or excessive missing data were excluded.

### 2. Reconstructing of mutation spectrum

Reconstruction of mitochondrial mutational spectra was performed using the NeMu pipeline, a framework specifically designed to infer mutation spectra from protein-coding sequences while accounting for phylogenetic structure and substitution context. NeMu integrates multiple steps including sequence alignment, phylogenetic tree inference, ancestral state reconstruction, and classification of nucleotide substitutions into standardized mutation categories. In this study, the pipeline was applied to mitochondrial protein-coding genes, and only synonymous substitutions were retained for downstream analyses to reduce the influence of selection at the amino-acid level.

Phylogenetic trees required for mutation inference were reconstructed using maximum-likelihood methods as implemented in IQ-TREE, with appropriate substitution models selected automatically. Ancestral states at internal nodes were inferred along the resulting trees, allowing mutations to be mapped onto individual branches. For each species, inferred substitutions were aggregated and summarized into 12-class mutational spectra, representing relative frequencies of all possible single-nucleotide substitutions.

All downstream data handling, visualization, and statistical analyses were conducted in Python, using standard scientific libraries for data manipulation (pandas, numpy), distance calculations (scipy), statistical testing (scipy.stats), and plotting. Cosine distance metrics were used to quantify similarity between mutational spectra, and both parametric and non-parametric statistical tests were applied to compare mutation frequencies between groups.

Custom scripts were used to integrate outputs from different steps of the pipeline, perform filtering based on mutation counts, and generate figures and summary tables. All analyses were conducted using reproducible workflows, ensuring consistency across datasets.

### 3. DNA metrics - GhAhSkew

The nucleotide composition of mitochondrial DNA was characterized using several metrics proposed in previous studies. All metrics were calculated from nucleotide frequencies at fourfold degenerate codon positions. Fourfold degenerate sites are a subset of third codon positions at which any nucleotide substitution does not alter the encoded amino acid. Because substitutions at these sites are largely free from selective constraints, they are commonly used as proxies for neutral evolution. Consequently, the nucleotide composition at fourfold degenerate positions is expected to be determined predominantly by the long-term mutational process. In this study, all metrics were calculated separately for each protein-coding mitochondrial gene in termites and cockroaches.

To characterize the balance between complementary nucleotide pairs, we used the GhAhSkew.

The GhAhSkew metric quantifies the relative excess of guanine over adenine at fourfold degenerate positions on the mitochondrial heavy strand. Because oxidative damage to mtDNA preferentially increases the probability of A→G substitutions over evolutionary time, GhAhSkew has been proposed as an indirect indicator of oxidative mutational pressure acting on the heavy strand. Thus, higher GhAhSkew values correspond to a greater relative accumulation of guanine compared with adenine, consistent with stronger long-term oxidative mutational bias. The metric was calculated as follows (Equation 1.1):

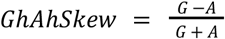

where *G* is the number of guanine nucleotides at fourfold degenerate positions on the mitochondrial heavy strand, and *A* is the corresponding number of adenine nucleotides.

### 4. Codon usage and synonymous codon usage bias

Codon usage was analyzed according to the vertebrate mitochondrial genetic code (NCBI translation table 2). Because mitochondrial protein-coding genes are encoded on both the majority and minority DNA strands and are therefore exposed to strand-specific mutational pressures, all codon usage analyses were performed separately for genes located on each strand. The minority-strand dataset included ND1, ND4, ND4L, and ND5, whereas the majority-strand dataset comprised ATP6 (A6), ATP8 (A8), CO1, CO2, CO3, CYTB, ND2, ND3, and ND6.

To evaluate differences in synonymous codon usage between termites and cockroaches, a species-level index of synonymous codon preference was calculated.

For each species, only twofold degenerate codon pairs differing exclusively at the third codon position were considered. For genes encoded on the majority strand, codon pairs ending in T and C were analyzed, whereas for genes encoded on the minority strand, codon pairs ending in A and G were used.

To eliminate the effect of amino acid composition, codon frequencies were normalized within each synonymous codon pair. Specifically, for majority-strand genes the normalized frequency was calculated as

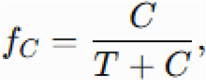

where *C* and *T* denote the genomic frequencies of codons ending in C and T, respectively. Similarly, for minority-strand genes,

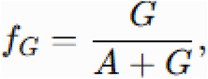

where *G* and *A* denote the genomic frequencies of codons ending in G and A, respectively.

Thus, each value represents the relative preference for one synonymous codon over its alternative and is independent of amino acid abundance.

For each species, the median normalized codon frequency was calculated across all synonymous codon pairs.

### 5. Prediction of amino acid composition changes from mutational spectra

To estimate the expected contribution of mutational spectrum differences to amino acid composition divergence, a codon-based probabilistic model was applied.

For each group independently, the expected changes in amino acid composition were predicted from the corresponding empirical mutational spectrum. The mutational spectrum was represented as the relative probabilities of the 12 possible nucleotide substitutions (e.g., A→G, C→T). For majority-strand genes, the mutational spectrum estimated from the CO1 gene was used directly. For minority-strand genes, the complementary mutational spectrum was applied, accounting for the fact that these genes are encoded on the opposite DNA strand.

For each mutation type mmm, the difference between termites and cockroaches was first calculated as

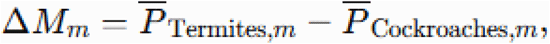

where P denotes the mean probability of the corresponding nucleotide substitution in the empirical mutational spectrum.

For each codon, all possible single-nucleotide substitutions were simulated across the three codon positions. The probability of each substitution was determined by the corresponding nucleotide substitution probability in the empirical mutational spectrum.

Each possible codon transition was converted into an amino acid transition using the mitochondrial genetic code. The contribution of each codon mutation to amino acid composition change was calculated as:

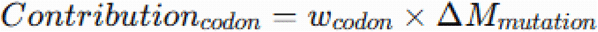

where wcodon represents the relative weight of the codon and ΔMmutation represents the probability of the corresponding nucleotide substitution.

In the current model, all coding codons were considered equally probable:

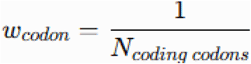

For each amino acid, all incoming and outgoing mutation flows were summed. The predicted change in amino acid composition was calculated as:

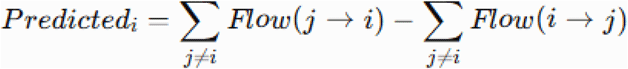

where the first term represents all mutational transitions generating amino acid *i*, whereas the second term represents all transitions resulting in the loss of amino acid *i*. Positive values indicate an expected increase in the abundance of amino acid *i*, whereas negative values indicate an expected decrease under the given mutational spectrum.

The procedure was performed independently for termite and cockroach mutational spectra, resulting in four predicted amino acid compositions: majority strand — termites; majority strand — cockroaches; minority strand — termites; minority strand — cockroaches.

The predicted difference between termite and cockroach amino acid compositions was subsequently calculated as:

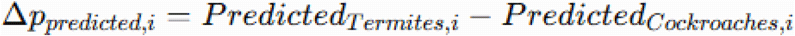

### 6. Divergence of amino acid composition and its correspondence to differences in the mutational spectrum

To quantify differences in amino acid composition between termites and cockroaches and to estimate the proportion of these differences consistent with divergence in mutational spectra, observed and predicted changes in amino acid composition were compared separately for the majority and minority strands.

For each amino acid *i*, the observed difference in amino acid composition was calculated as the difference between its mean relative abundance in termites and cockroaches:

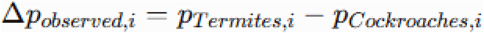

where pTermites,i and pCockroaches,i represent the mean relative frequencies of amino acid *i* in the corresponding groups.

An amino acid change was considered consistent with the mutational spectrum when observed and predicted differences had the same direction:

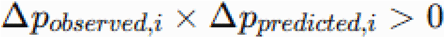

Thus, an amino acid was classified as consistent when its abundance was higher in termites than in cockroaches in both the observed and predicted compositions, or when its abundance was lower in termites in both cases.

The total magnitude of amino acid composition divergence between termites and cockroaches was quantified as the sum of absolute observed differences:

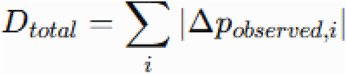

This metric represents the overall magnitude of amino acid composition divergence and weights each amino acid according to the size of its observed difference.

The divergence component consistent with mutational spectrum predictions was calculated as:

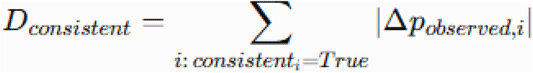

The proportion of total amino acid composition divergence consistent with differences in mutational spectra was calculated as:

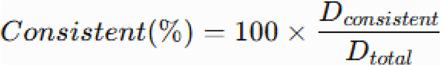

## ACKNOWLEDGMENTS

We are very grateful to Dr Alina Mikhailova for guiding and mentoring this work.

